# NETSseq Reveals Inflammatory and Aging Mechanisms in Distinct Cell Types Driving Cerebellar Decline in Ataxia Telangiectasia

**DOI:** 10.1101/2025.05.13.653662

**Authors:** Stirparo G Giuliano, Xu Xiao, Thompson Toni, Page Keith, Harvey RM Jenna, Cadwalladr David, Lawrence Jason, Burley J Russell, Juvvanapudi Joel, Roberts Megan, Barker F Daniel, Mulligan Victoria, Sherlock Chloe, Lizio Marina, Christie Louisa, Mudaliar Mani, Sheardown Steven, Margus Brad, Thompson Craig, Dickson Louise, Brice L Nicola, Carlton B Mark, Powell AC Justin, Dawson A Lee

## Abstract

The cellular and molecular changes driving the neurological abnormalities associated with ataxia – telangiectasia (A-T) are not well understood. Here, we applied our proprietary Nuclear Enriched Transcript Sort sequencing (NETSseq) platform to investigate changes in cell type composition and gene expression in human cerebellar post-mortem tissue from A-T and control donors. Compared to single-nuclei technologies, NETSseq provided a more robust detection of genes with low abundance, a higher cell type specific expression pattern, and significantly lower levels of cross-contamination. We found dysregulation in neurotransmitter signaling in granule neurons, potentially underlying the impaired motor coordination in A-T. Astrocytes and microglia have evidence of accelerated aging, with astrocytes being characterized by neurotoxic signatures, while microglia showed activation of DNA damage response pathways. These findings highlight the importance of NETSseq as a resource for investigating mechanisms and biological processes associated with disease, providing high-sensitivity, cell-specific insights to advance targeted therapies for neurodegenerative diseases.

## Introduction

Ataxia–telangiectasia (A-T) is a rare, autosomal recessive, multisystem disorder caused by mutations in the Ataxia–Telangiectasia Mutated (ATM) gene. A-T is characterized by a devastating and progressive neurological pathology. The first symptoms manifest in the early years of life as ataxia and loss of motor coordination, and later in life with oculomotor apraxia, dysarthria, choreoathetosis, drooling and difficulties swallowing. Loss of ATM protein function leads to a range of symptoms, including susceptibility to neoplasia, immunodeficiency resulting in sinopulmonary infections, and extreme sensitivity to ionizing radiation. Individuals with A-T are usually wheelchair-bound by their teenage years and the disease is often fatal in the second or third decade of life.

ATM is a member of the serine/threonine protein kinase family^1^ that has homology with a family of phosphatidyl inositol 3’ kinase-related kinases (PIKKs), a group of high molecular mass eukaryotic proteins found in all cells. ATM protein is primarily activated by double stranded breaks (DSBs) in DNA and regulates the cellular response to DNA damage, including DNA repair mechanisms, replication checkpoints, and other metabolic changes. Accumulating DNA damage can cause cells to exit the cell cycle and become senescent or undergo programmed cell death to restrict the propagation of mutational events^2^. Although ATM is central to this signaling cascade, there is a highly coordinated series of molecular events involving several other key factors which converge when the cell detects DNA damage. Interestingly many of these factors are also substrates of and thus regulated by ATM^3^. ATM has also been shown to play diverse roles in cellular metabolism, mitochondrial redox sensing, regulation of antioxidant capacity, and potentially autophagy and lysosomal trafficking^4,5^.

Despite decades of research on ATM^6^ and its role in responding to DNA damage^7,8^, less is known about how ATM might function in different cellular contexts, and whether these differing functions might account for the pleiotropic symptoms seen in A-T patients. One of the earliest symptoms of A-T is ataxia, or the impairment of motor coordination, which results from progressive degeneration of the cerebellum likely resulting from a progressive loss of both Purkinje and granule neurons^9^. While numerous potential mediators of this neurodegeneration have been proposed, the precise cause is likely to be a complex interaction between diverse cellular mechanisms that impact overall neuronal health and survival. This may include not just the deficient response to DNA damage, but also impaired glial support, mitochondrial dysfunction, altered oxidative stress and inflammatory responses^10^, all ultimately leading to deficits in neuronal and synaptic function. Since there is currently no cure for A-T and no therapeutics that alter the disease progression, treatments are limited to partial symptomatic relief. For this reason, decoding the cell type specific response to DNA-damage-induced cellular pathology is vital for developing a cohesive biological strategy in the pursuit of new therapeutic interventions.

To obtain deep, cell type specific transcription profiles from the cerebellum of patients with A-T and control donors, we used Nuclear Enriched Transcript Sort sequencing (NETSseq)^11^, which combines fluorescent activated nuclei sorting (FANS) to purify nuclei from cell types of interest with RNA-seq to generate genome-wide transcriptional profiles. Using NETSseq, we generated 318 RNA-seq samples (248 control and 70 A-T), covering eight distinct cerebellar cell types – granule, basket, Golgi and Purkinje neurons, as well as astrocyte, microglia, mature oligodendrocyte, and oligodendrocyte precursor (OPC) glial cell types. Compared to single cell technology, NETSseq offers more robust detection of genes with low expression and better captures differential expression across cell types and disease. We found that disease-associated gene expression changes were remarkably cell type specific, offering novel insights into the differing mechanisms within these cell types that contribute to neurodegeneration and motor dysfunction in A-T.

## Results

### NETSseq: Isolation of Cell type specific Nuclei

Cerebellar post-mortem tissue samples from 15 donors with a clinical diagnosis of A-T and 56 control donors were used in these analyses. The A-T cohort comprised five female (8-31 years old) and ten male (8-32 years old) donors with neuropathological findings consistent with the disease. Control donors (27 males aged 8-93 years and 29 females aged 17-96) were selected for having no known CNS disease and having died from a non-CNS related etiology. Post-mortem intervals ranged from 2-21 h for A-T donors and 5-79 h for controls donors. Donor information is summarized in Table S1.

To better understand the cell type specific molecular changes that occur in A-T, we extended the NETSseq cell type purification method from Xu et al.^11^ to isolate additional cell types from post-mortem human cerebellar tissue (Fig. 1A). Using a combination of cell type specific antibodies, we developed FANS gating strategies to isolate specific populations of nuclei and determine the relative abundance of each cell type (Fig.1B, Table S2). Using a refined antibody panel, we were able to isolate nuclei from an additional three key cerebellar cell types – microglia, Golgi and Purkinje neurons, in addition to the five cell types isolated in the previous publication (granule, basket, astrocyte, oligodendrocyte and OPCs). To isolate the four neuronal cell types, we used antibodies against the splicing factor NeuN/RBFOX3, the ER resident membrane protein Inositol 1,4,5-Triphosphate Receptor Type 1 (ITPR1), and the transcription factor Forkhead Box K2 (FOXK2). Combining ITPR1 and FOXK2 labeling into a single channel and plotting this against NeuN in a different channel, enabled efficient detection of pure populations of nuclei from granule and basket neurons, along with those from exceedingly rare Golgi (0.1%) and Purkinje (0.01%) neurons. The complete sorting strategy for Purkinje neurons is summarized in Fig.S1D. We also isolated oligodendrocytes and OPCs based on differential positive labeling of oligodendrocyte transcription factor 2 (OLIG2), and astrocytes based on positive labeling of Glutamate Aspartate Transporter (GLAST/EAAT1/SLC1A3). Finally, we confirmed that labelling against interferon regulatory factor 8 (IRF8), which we previously used to isolate cortical microglia^12^, is also effective for isolating cerebellar microglia.

**Fig. 1.**
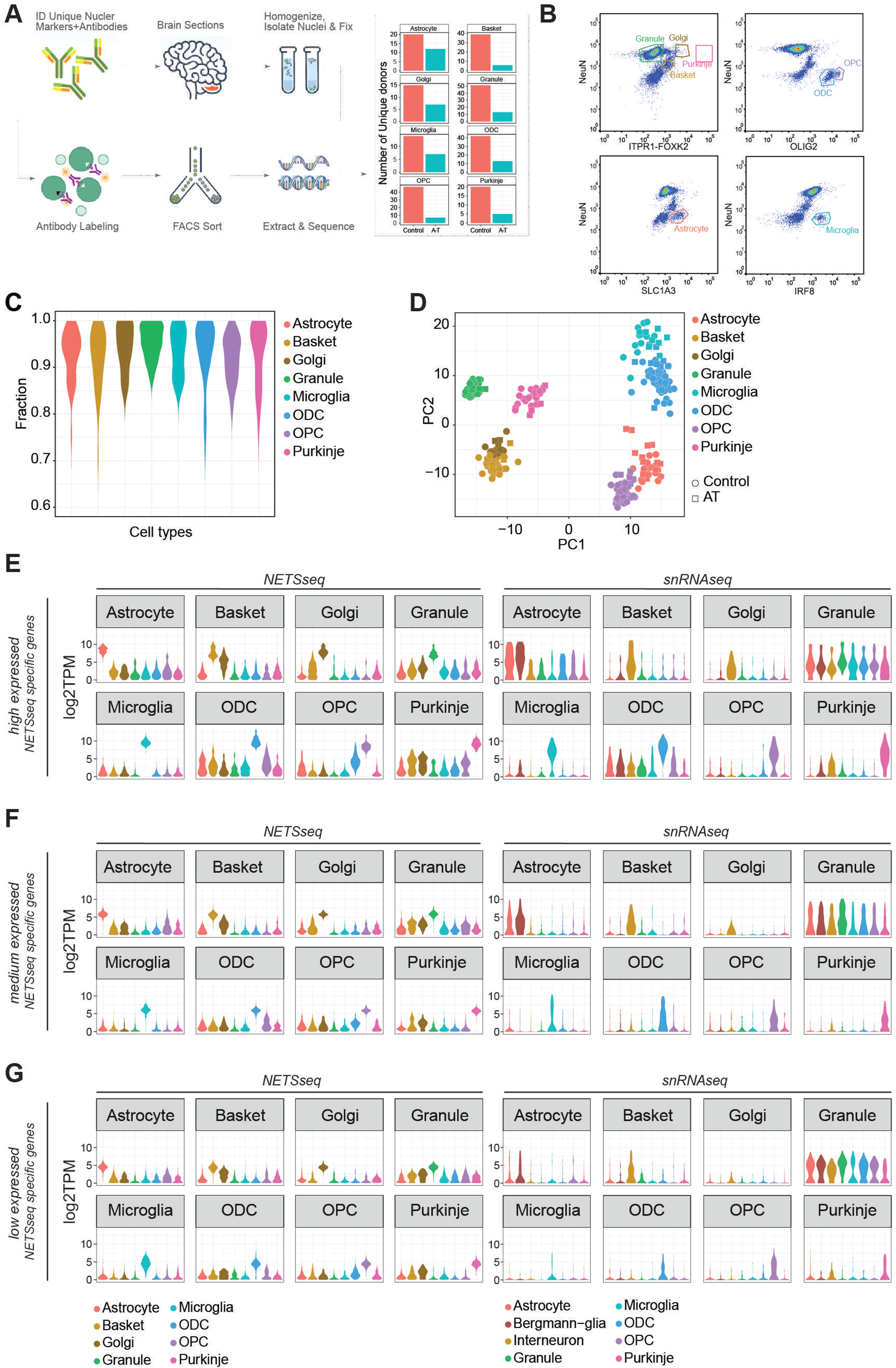
(A) Schematic flow of our Nuclear Enriched Transcript Sort sequencing (NETSseq) pipeline and summary barplot of the donors and cell types analysed in this study. (B) Representative FACS plots showing populations of stained nuclei and the gating strategy. Antibodies used are noted against the axes of the plots. Gatings for relevant populations are highlighted and the cell types are annotated. (C) Distribution of fraction of similarities across cell types indicates clean populations. Fractions were computed using the DeconRNAseq R package. (D) Principal component analysis of the cerebellum cell types, computed using the top 500 variable genes, shows tight and clean clusters. (E,F,G) Violin plots showing the expression in both NETSseq and pseudobulked snRNA-seq data sets of cell type specific genes calculated from the NETSseq results. The specific genes were identified in the NETSseq cohort after filtering for (E) high expressed genes (>66% of expression quantile), (F) medium expressed genes (33%-66% of expression quantile) and (G) low expressed genes (<33% of expression quantile).

### Deep Expression Profiling of the Human Cerebellum

RNA-seq was used to generate nuclear gene expression profiles for each population of nuclei separated by FANS. The number of nuclei collected per donor varied by cell type from a mean of 1,096 for Purkinje neurons to 90,720 for granule neurons. From each sorted nuclei population, we subsequently generated deep RNA-seq libraries (median reads >16 million after adapter trimming, Table S1). To validate the identity and purity of nuclei obtained from each sorting strategy, we evaluated the expression of known markers for each cell type (Figure S1A) and performed a deconvolution analysis (Fig.1C). This involved assembly of a reference signature for each cell type; then, for each sample, we estimate the fraction of similarity against all the cell types in the assembled reference. Consistent with previous findings^11^ each cell type exhibited a specific expression pattern of marker genes, indicating that negligible contamination was present between cell types.

Next, we used two unbiased methods, hierarchical clustering (HC, Fig.S1B) and principal component analysis (PCA, Fig.1D), to visualize the similarities and differences across samples. With both methods, we observed a high level of similarity for samples from the same cell type and clear separation of samples from different cell types. Using these methods, we can also observe similarities between basket and Golgi cells, which are two subtypes of GABAergic interneurons. Cell type specific expression of key cell type specific marker genes was confirmed using *in situ* hybridization (ISH) analysis in human donor tissue (Fig.S1C). These results indicate that NETSseq can purify and profile distinct cell types with high accuracy and low levels of cross contamination.

### NETSseq Technology Provides Enhanced Capability for Detecting Low Expressed Cell Type Specific Genes

Single-nuclei analysis provides superior resolution and the capacity to delineate cellular heterogeneity compared to bulk analysis; however, it faces limitations in detecting genes with low expression profiles. To assess the detection thresholds and sensitivity of our NETSseq platform against single-cell technologies, we performed a comparative analysis, directly comparing our findings with those of Lai et al.^13^, which generated a snRNA-seq dataset from the cerebellum of control and A-T donors. First, we identified cell type specific genes in both datasets and categorized them based on expression quantiles (low: <33%, medium: 33%-66%, high: >66%). We generated expression distribution plots (Fig.1E-G, Fig.S3A) and heatmaps (Fig.S2A-D) for these genes across all examined cell types in both datasets. In all snRNA-seq expression tertiles, computed using the specific genes generated by NETSseq, we noted the presence of granule neuron markers in the single cell data for the non-granule cell types. This may be due to the presence of ambient RNA^14,15^ originating from the high number of granule neurons, which comprise ∼70-90% of total cells. This contamination was also found when the expression of snRNA-seq-derived specific genes were plotted across all cell types in the snRNA-seq study (Fig.S3A). Moreover, for the lower tertiles, the distribution of gene expression for the specific genes in each cell type in the single-cell cohort is significantly skewed toward zero, indicating limited sensitivity in detecting these genes (Fig.1G). In contrast, when we examined the specific genes identified in each expression tertile in the single-cell cohort, we observed no loss of signal in the NETSseq dataset (Fig.S2C); this suggested that NETSseq did not face such limitations and was able to detect the majority of the specific genes.

To determine whether the NETSseq gating strategy for each cell type accurately isolated the target cell type, we performed a topological analysis using UMAP generated from the snRNA-seq data, by comparing the overlap between the snRNA-seq assigned cell types and the snRNA-seq derived expression of the top cell type markers identified from NETSseq data (within the highest expression quantile). The analysis revealed a complete overlap between the specific markers and the regions covered by individual cell types, confirming that the correct populations were sorted, without contamination from other cell types (Fig. S3B). Furthermore, this analysis demonstrated that the sorted populations for each cell type capture the entire cell type population, without any biases for any subpopulations. The only exception is cerebellar astrocytes, which can be split into three regionally, morphologically, and molecularly distinct subpopulations i.e. specialized Bergmann glia (BG), which are intimately connected to the Purkinje cell bodies, granule layer and white matter astrocytes. snRNA-seq data from Lai et al. subdivided astrocytes into two subgroups i.e. BG and other astrocytes. Comparison of expression profiles shows that our sorted astrocytes contain both BG and other astrocytes (Fig. 1E-G, Fig. S2C).

### Cerebellar Cell Type Composition in Control Donors

Histological studies have shown a progressive loss of Purkinje neurons and to a lesser degree, granule neurons in A-T patients^10,16,17^. However, less information is available regarding the proportions of other cerebellar cell types and how these change with disease progression. To address these questions, we used various methods to characterize the typical cell type composition of the cerebellum, and to examine how cell type proportions change in disease.

First, we analyzed the flow cytometry profiles obtained during nuclei sorting to quantify the relative proportion of each cell type population (Table S2). Consistent with previous stereology studies^18,19^, we found that granule neurons accounted for 81-93% of the cells in cerebellum, whereas Purkinje neurons are extraordinarily rare, comprising 0.001-0.027% of the cells. Due to the abundance of granule neurons, we found that cell types such as astrocytes and mature oligodendrocytes, which are normally abundant in other brain regions, are relatively rare in the cerebellum, comprising only 2.2% and 1.3% of cells, respectively. To complement this analysis, we estimated cell type proportions using global gene expression deconvolution (Fig.2A). Briefly, for each cerebellar sample we performed RNA-seq on unsorted nuclei, which could then be deconvoluted using reference signatures from each constituent cell type, to obtain cell type proportions of the different cell types. The deconvolution analysis yielded similar results to those from the flow analysis, showing granule neurons as the predominant cell type at 70%, while Purkinje neurons were the least common at 0.7%. Both methods have technical limitations: flow analysis, limited by the efficiency of antibody labeling, likely underestimates cell type proportions, while deconvolution, biased by nuclear size and RNA content, tends to underestimate the proportions of cell types with smaller nuclei (such as granule neurons and glia) and overestimate those with larger nuclei (Purkinje, Golgi, basket neurons). Importantly, the two methods provide relatively consistent proportions and align with cell type proportions determined using stereology^18,19^ and single nuclei RNA-seq methods^13^ (Fig.2A, Fig.S3C).

**Fig. 2.**
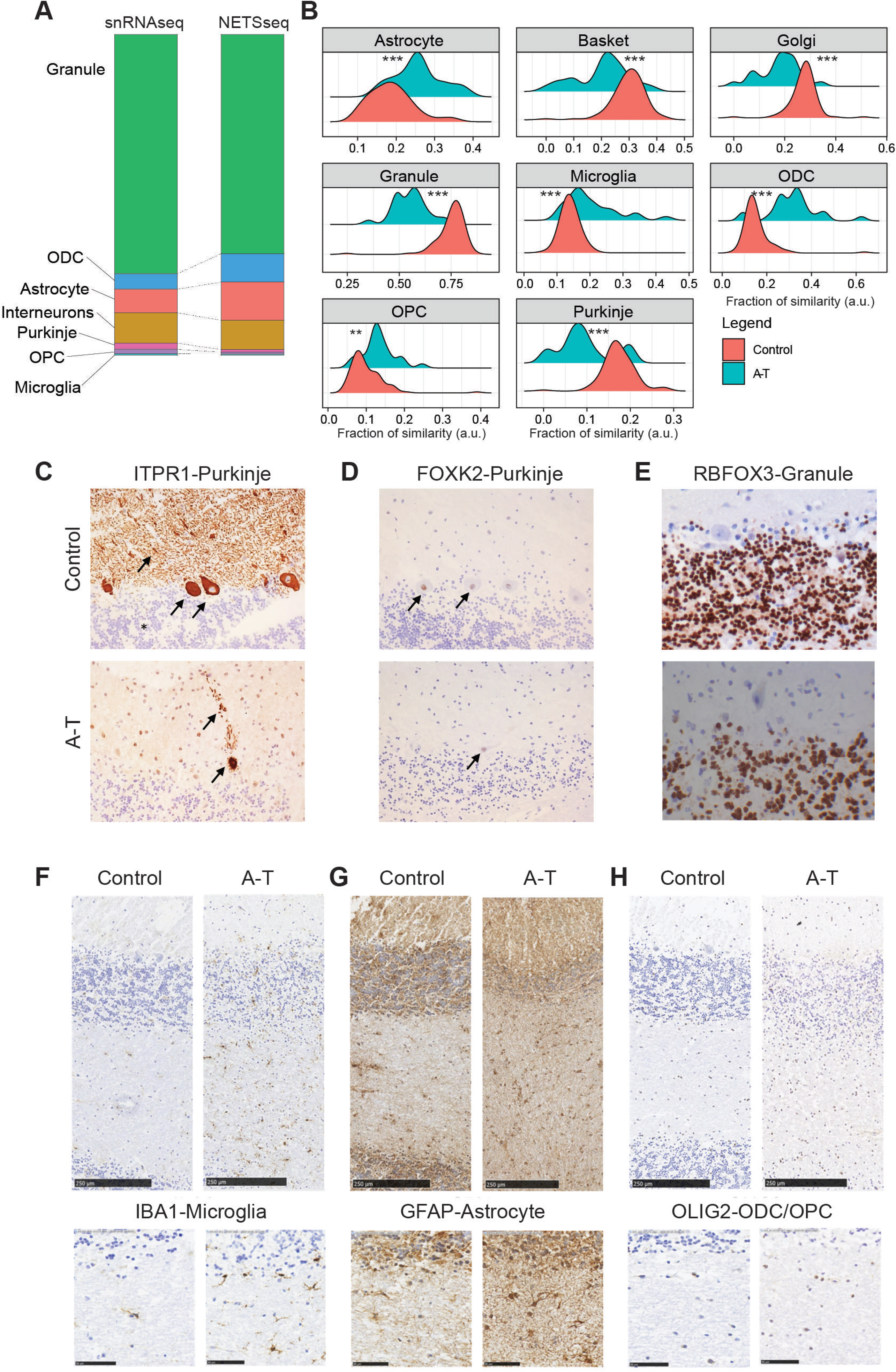
(A) Stacked bar plot showing the similarities of cell type proportion computed using the snRNA-seq and NETSseq cohorts. (B) Relative fraction of similarity between A-T and control donors across the examined cell types. The fractions represent the similarity of unsorted samples, stratified by disease condition, to reference signatures constructed for each cell type. Each reference signature comprises the expression profile of the target cell type and the average expression of all remaining cell types (Control N=100; A-T N=25, pairwise t.test with BH for multiple comparison). (C) Photomicrographs (20×) showing representative patterns of ITPR1. Arrows show cytoplasm and dendrites of Purkinje cells, asterisk indicates Granule cells. (D) FOXK2 immunoreactivity in one healthy control (donor MD_5669) and one A-T (MD_5902) donor. Arrow indicates Purkinje neuron nuclei. (E) RBFOX3 staining in granule neurons from both donors, expression was not observed in Purkinje cells (40×). (F-H) IHC showing glial cell markers in sections of cerebellar cortex from one healthy control donor (MD_5751), and one A-T case (MD_5902). Top row shows sections extending from the molecular layer through the granule cell layer and into the white matter (scale bar is 250 µm). Bottom row shows details of the white matter (scale bar is 50 µm). Antibodies for IBA1 identify microglial cells (F), GFAP for astrocytes (G) and OLIG2 for ODCs and OPCs (H). Relative to control tissues, glia were more numerous in A-T donor sections.

### Changes in Cerebellar Cell Type Proportions in A-T

Next, we aimed to identify changes in cell type proportions in A-T. Given the imbalance of cell type proportions in the cerebellum, where most cell types skew towards zero, we employed a modified deconvolution approach which more accurately captures differences between two conditions. Here, rather than conducting a global deconvolution, where an unsorted sample is split into all constituent cell types and the proportion of these cell types must add up to 1, we used a comparative deconvolution approach. This method assesses a single cell type at a time, making it more useful for measuring changes across conditions rather than absolute cell type proportions.

The density distribution of the relative abundance for each cell type shows the change of cell composition between A-T versus controls (Fig.2B). We detected a significant (t.test with Benjamini-Hochberg correction for multiple comparison) loss of Purkinje neurons, with a decrease of 1.85-fold, which aligns with the typical cerebellar degeneration seen in ataxia^20^. This degeneration contributes to the impaired motor coordination and balance deficits experienced by affected individuals. We also observed a significant loss of granule neurons with a reduction of 1.35-fold (Fig.2B, Fig.S3C), highlighting the extensive neuronal loss within the cerebellum. This loss of granule neurons further exacerbates the dysfunction of cerebellar circuits, leading to the pronounced clinical symptoms of ataxia. Consistent with more general neuronal loss, we also observed a decrease of Golgi and basket neuronal cell types with a reduction of 1.52-fold and 1.45-fold, respectively. Additionally, our analysis revealed a 1.35-fold increase in OPCs and a 2-fold increase in ODCs, (Fig.2B, Fig.S3C, Table S3), suggesting a compensatory response to ongoing myelin damage^21^. Astrocytes and microglia show a 1.37- and 1.44-fold increase, respectively, in cell numbers.

We further confirmed the decrease in granule neurons and increase in mature oligodendrocyte numbers using flow analysis. Because the fluorescence profiles for these two cell types were distinct and well separated from other populations in both control and A-T donors, we could confidently quantify their proportions across a subset of donors. This analysis showed that almost half of granule neurons degenerate in A-T, decreasing from around 90% in control to 55% in A-T donors. In contrast, oligodendrocytes increase in relative proportion by approximately 4-fold, from 1.5% in control donors to 6% in A-T donors (Fig. S3D).

### Histopathological Evaluation of Neuron Loss and Glial Activation in A-T Cerebellum

On a subset of cerebellar cortical tissue samples, we performed histopathological analysis to evaluate gross neuropathological features in control and A-T donors (Fig.2C-H). In this analysis, we used antibodies against ITPR1 and FOXK2 to label Purkinje neurons (Fig.2C,D), RBFOX3 to label granule neurons (Fig.2E), IBA1 for microglia (Fig.2F), GFAP for astrocytes (Fig.2G), and OLIG2 for oligodendrocytes and OPCs (Fig. 2H). In control Purkinje neurons, FOXK2 is normally localized to the nucleus, while ITPR1 labels the cell body, including dendritic arborizations. Qualitative analysis in A-T donors showed a clear reduction in Purkinje neuron number (Fig.2C,D), while surviving Purkinje neurons have features characteristic of neurodegeneration, including disturbed morphology and ectopic appearance of dendrites in the molecular layer (Fig.2C). Importantly, although Purkinje neurons in A-T donors are reduced in numbers, they can still be reliably labelled using antibodies against both ITPR1 and FOXK2. Qualitative analysis of glial cell markers in A-T donors showed more abundant IBA1 microglia labelling (Fig.2F, Fig.S3E), more abundant and more intense GFAP labelling in astrocytes (Fig.2G, Fig.S3E) and more abundant OLIG2 oligodendrocyte lineage labelling, particularly in the white matter (Fig.2H).

Finally, for granule neurons, we performed quantitative analysis using tissue sections from three control and four A-T donors. We observed a 70% decrease in granule neuron numbers in A-T donors, which is consistent with the approximately 50% reduction from the comparative deconvolution analysis. Overall, the histology results support the finding that neuronal cell types decrease, while glial cell types increase in the disease. Furthermore, the increased expression of GFAP suggests increased astrocyte activation in A-T donors.

### Distinct Cell type specific Gene Expression Signatures in NETSseq Samples

We conducted a comparative analysis of gene expression values to assess the similarities between our cohort and the single-cell cohort, by computing the pairwise correlation across each cell type (Fig.3A). Interestingly, we observed a strong overall correlation between the gene expression signatures of corresponding cell types across the two cohorts (Fig.3A); however, we detected granule neuron contamination in all cell types in the snRNA-seq dataset (Fig.3B). Moreover, our expressed genes demonstrate greater specificity to individual cell types compared to those in the single-cell cohort (Fig.3B).

**Fig. 3.**
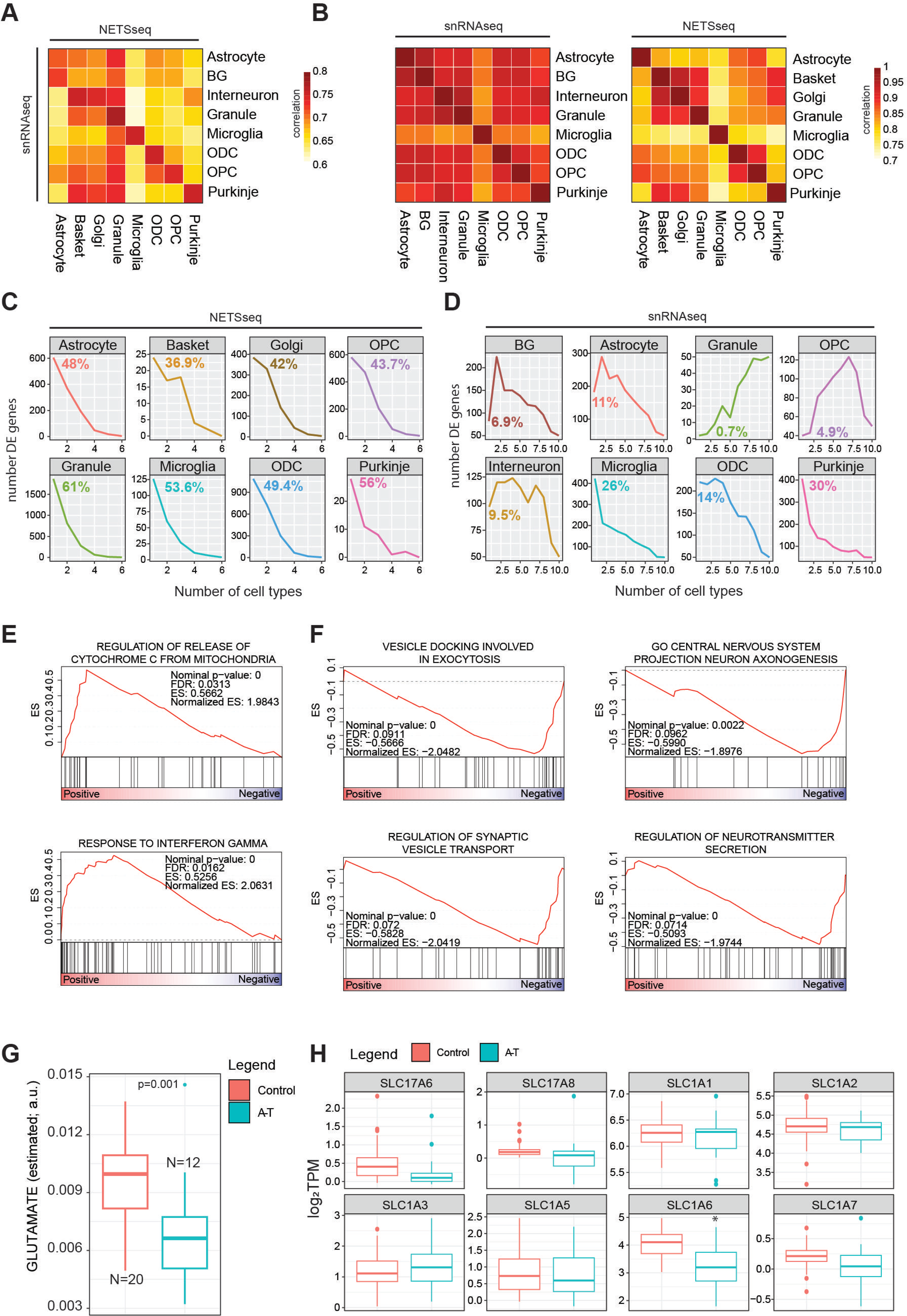
(A) Gene expression correlation analysis across cell type between the NETSseq and snRNA-seq cohorts and (B) within each cohort. (C) Line plot showing the occurrence of differentially expressed genes across cell types in the NETSseq cohort and (D) single nuclei cohort. For each plot we highlighted the percentage of specific differentially expressed genes in each cell type. (E) GSEA results for the upregulated differentially expressed genes between A-T and control in granule cells. (F) GSEA results for the downregulated differentially expressed genes between A-T and control in granule cells. (G) Distribution of glutamate metabolite levels, estimated with Metabolic Flux Analysis using the scFEA R package (see methods), in granule neurons. The significance was derived using the scFEA R package, the y axis represents the normalized amount of estimated glutamate (a.u.: arbitrary unit) (H) Expression boxplot derived from the NETSseq cohort shows a significant downregulation of the SLC1A6 transporter; the significance was computed using the DEseq R package (Control=20, A-T=12, p.adj<0.05).

We next performed differential gene expression analysis comparing A-T and control samples across all cell types (Fig.S4A, Table S4). The presence of technical variance was accounted for and corrected in the DESeq model using surrogate variables (Fig.S4B). Moreover, we evaluated if any correlation existed between the number of nuclei collected and the transcriptomic signal and found no significant correlation between these two variables (Fig. S4C). To validate our findings, we compared our results with those reported in the snRNA-seq study by Lai et al^13^ (Table S5). Intriguingly, in our cohort, we observed that a higher proportion of differentially expressed genes are cell type specific (Fig.3C). In contrast, this trend is not uniformly observed in the study of Lai et al., particularly in granule neurons where differentially expressed genes overlap more extensively with other datasets (Fig.3D). These results highlight the ability of NETSseq to resolve these cell type specific expression changes, particularly for granule cells (Fig.3C), where the differences in the snRNA-seq data are likely obscured by ambient RNA contamination across all cell types (Fig.3D).

### Granule Neuron Dysfunction and Neurotransmission Imbalances

In the cerebellar cortex, mossy fibres (MF) initiate a cascade where they stimulate granule neurons, which subsequently activate Purkinje neurons. Together, these signals are essential for motor coordination^22^, and disruptions in Purkinje neurons are known to lead to motor impairments such as ataxia. In their study, Joon-Hyuk et al.^22^ emphasized the critical role of granule neurons in motor function by showing that the integrity of MF-induced signaling and consistent firing of Purkinje neurons relies on granule neuron function. Lai et al.^13^ observed a significant loss of granule neurons in A-T; however, challenges in single-cell technology prevented the detection of differentially expressed genes. Our NETSseq platform can overcome those limitations and better elucidate the role of granule neurons in the cerebellum of A-T donors. GSEA analysis performed with the upregulated genes in granule neurons in A-T vs control donors, highlighted the gene set "Release of cytochrome C from mitochondria". This process is crucial for apoptosis regulation and is influenced by interferon gamma (IFN-γ), a “response to interferon gamma” gene set was also significantly enriched (Fig.3E). We also noted a significant downregulation of BCL2, an anti-apoptotic protein, in A-T donors, suggesting compromised regulation of apoptosis (Table S4). Downregulation of synaptic vesicle transport in cerebellar granule neurons may disrupt neurotransmitter release, impairing excitability of Purkinje neurons and interneurons^23^. Consistent with this observation, processes associated with neurotransmitter release and vesicle docking were markedly enriched among downregulated genes in A-T granule neurons, indicating potential disruptions in synaptic function associated with motor coordination mechanisms (Fig.3F). Metabolic flux analysis computed using NETSseq gene expression values aggregated across entire pathways, suggested that granule neurons in A-T donors may have lower glutamate levels (Fig.3G), likely impairing synaptic transmission within the cerebellum and contributing to functional deficits. Specifically, we noticed a significant downregulation of the glutamate transporter SLC1A6 (Fig.3H), which is crucial for the clearance of glutamate from the synaptic cleft following synaptic release. By taking up glutamate into the cells, SLC1A6 helps maintain low extracellular glutamate concentrations, preventing excitotoxicity and ensuring precise synaptic signaling. Pathways associated with glutamate receptor signaling are also downregulated in Purkinje neurons of A-T donors, indicating a broader disruption of glutamatergic transmission throughout the cerebellum. The reduced glutamate receptor signaling in Purkinje neurons likely impairs neuronal function and exacerbates motor coordination deficits since these neurons play a central role in controlling cerebellar output (Fig.S4D). Taken together, the changes in glutamate levels and the dysregulated synaptic transmission processes may indicate a broader disruption in the balance / control between excitatory and inhibitory signaling. This dysregulation, together with the loss of granule neurons could impair the integrated timing and coordination of motor commands. These findings suggest that disruptions in glutamate neurotransmission contribute to the motor dysfunction observed in A-T, highlighting potential therapeutic targets for future studies.

### Expression Changes in Astrocytes are Consistent with Elevated Neurotoxic Activation and Inflammation in A-T

“Reactive astrocytes” refers to astrocytes that are undergoing morphological, molecular and functional remodeling in response to adverse microenvironmental changes and pathological stimuli. Astrocyte activation is implicated in the pathogenesis of multiple neurodegenerative diseases. Extensive changes were observed in our NETSseq derived astrocyte A-T datasets. Three mouse astrocyte activation gene expression signatures have previously been reported, corresponding to a common ‘pan’ activation profile, an ‘A1’ neurotoxic activation profile and an ‘A2’ neuroprotective profile^24^. Expression analysis indicated that the ‘A1’ neurotoxic and common “PAN” gene signatures are significantly upregulated in A-T donors, compared to controls (pval< 0.005) (Fig.4A), while the A2 signature was not significantly different. In addition to the genes specified in the A1 gene set, release of complement components is also characteristic of the A1 activation state and is postulated to drive synapse degeneration^24^. Our data shows that several activators of complement components are significantly upregulated in A-T donors compared to healthy controls (Fig. 4B). Conversely, most complement inhibitors have higher expression in controls, further supporting the hypothesis of an overall activation (Fig. S4E). Interestingly, CFI, a complement inhibitor, is significantly upregulated in A-T donors, possibly as a compensatory mechanism in response to the excessive complement activation^25^. While there are nuances and complexities related to astrocyte subset identification, function, and nomenclature of activation^26,27^, the findings mentioned above suggest an overall increased activation state in astrocytes in A-T. This heightened state may lead to an elevated neuroinflammatory response and burden, potentially contributing to the ongoing pathophysiology of the disease.

**Fig. 4.**
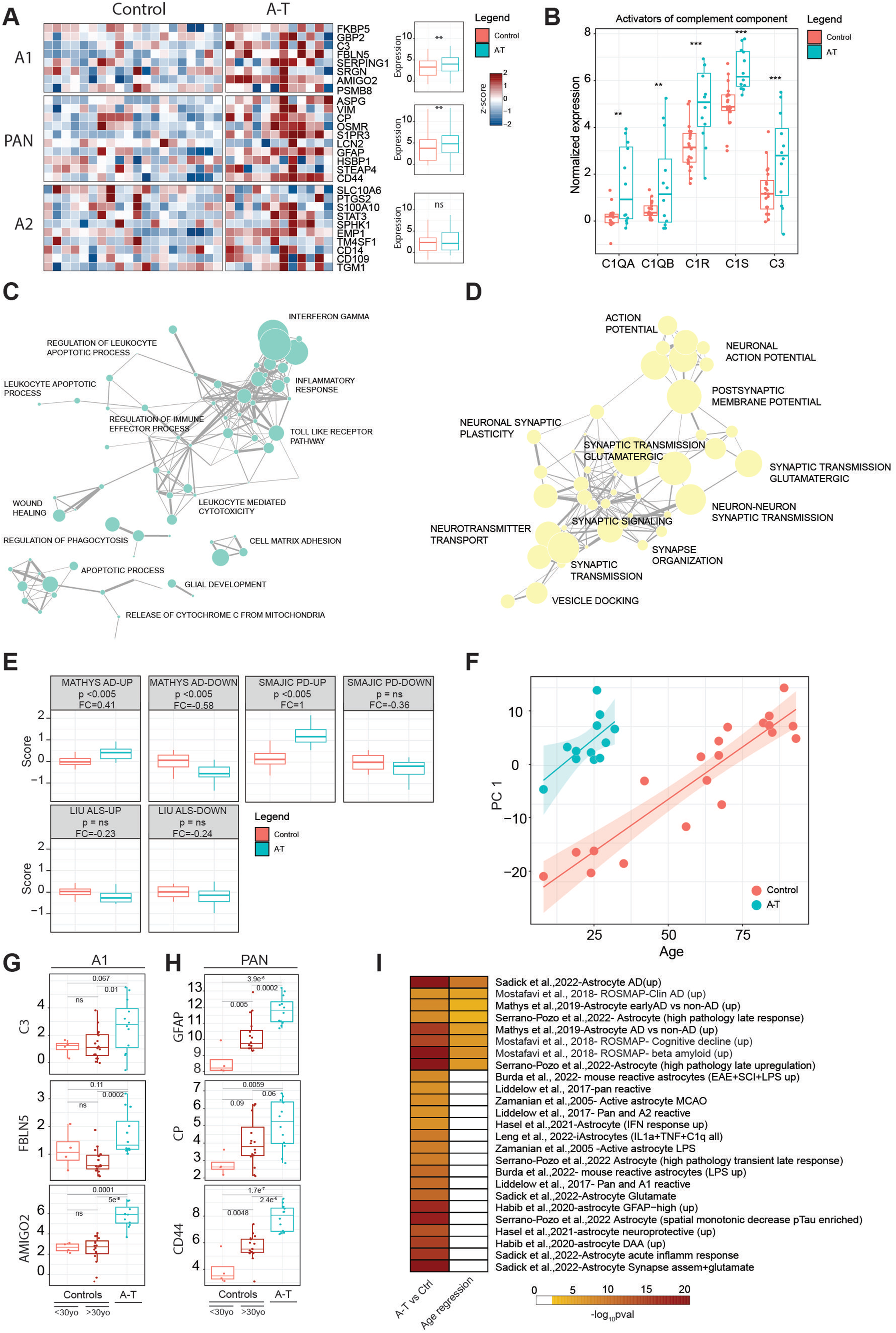
(A) Heatmap showing the expression of genes associated with A1, A2 and PAN signatures in A-T and control donors. For each signature, the expression values of individual genes in control or A-T donors were aggregated in a boxplot and the significance was computed using a pairwise t-test statistic with BH correction. (B) Expression of activators of the complement component in control or A-T donors in astrocyte samples. The significance was computed using a pairwise t-test statistic with BH correction. (C) Network of the most significant biological process enriched for the upregulated or (D) downregulated genes in Astrocytes. (E) Distribution of the gene set scores for signatures associated with changes in neurodegenerative disease in human AD^30^, PD^33^ and a mouse ALS model^32^ tested against expression changes observed in A-T or control astrocyte samples. p, t test, corrected for multiple comparison; Δ, log2 fold change in score. Gene sets are separated into up-regulated and down-regulated sets according to fold change in the source publication. (F) Scatter plot between PC1 dimension computed with the significant age-related genes in astrocytes and age of A-T or control donors. Linear regression analysis was split for disease condition. (G) Expression distribution of A1 associated markers (C3, FBLN5 and AMIGO2) in younger control donors (<30yo), older control donors (>30yo) and A-T donors in astrocytes. The significance was computed using a pairwise t-test statistic with BH correction. (G) Expression distribution of PAN associated markers (GFAP, CP and CD44) in younger control donors (<30yo), older control donors (>30yo) and A-T donors in astrocytes. The significance was computed using a pairwise t-test statistic with BH correction. (H) GSEA computed between the upregulated genes identified in A-T vs control analysis or using age as a variable of interest; gene sets are found in Table S7.

### Astrocytes Show Extensive Evidence of Synaptic Remodeling and Loss of Neurotransmitter Signaling Function

An important function of astrocytes in the healthy brain is to support synapse function, including uptake of neurotransmitters and release of trophic factors essential for neuronal survival and communication. As astrocytes become more reactive, they lose their capacity to support synapses and normal neuronal function^28^. Given that A-T is a neurodegenerative disease, we were particularly interested in exploring how genes involved in glial support of synaptic function were altered. To address this, we performed GSEA on the genes that were either upregulated or downregulated in astrocytes from donors with A-T. We then conducted a network analysis to identify and group together similar biological processes. Our findings revealed that upregulated genes in A-T astrocytes were predominantly associated with inflammatory processes, consistent with our earlier observation of an increased neurotoxic and activated glial signature. This suggests that in A-T, astrocytes may be shifting towards a more reactive and potentially harmful state (Fig.4C). Downregulated genes were involved in synaptic function and neuronal interaction (Fig.4D) indicating a very extensive disease driven remodelling and synaptic reorganization in the cerebella of donors with A-T. In addition, genes critical for maintaining the structural integrity of the astrocyte-synapse interaction, as well as those involved in monitoring the activity of adjacent synapses (such as neurotransmitter receptors, ion channels and transporters) were significantly downregulated. This downregulation extends to receptors for the primary neurotransmitters in the cerebellum, i.e. glutamate and GABA, both of which play essential roles in maintaining normal cerebellar function. The marked reduction in the expression of these receptors (Fig. S5A) indicates a significant impact of A-T on synaptic communication. The typical profile for our cerebellar astrocytes suggests a considerable similarity to that of Bergmann glia. These are highly specialized cerebellar astrocyte populations which have a close and highly interconnected interaction with Purkinje neurons, controlling and supporting the synaptic function of these rare neurons^29^. Thus, the observed deficits seen in their function could be much more significant in the cerebellum and long term have a greater impact on the disease progression.

### Astrocyte Changes Overlap with Those Seen in Other Neurodegenerative Diseases

Astrocyte specific expression profiles from post-mortem tissue from patients with Alzheimer’s disease (AD) have previously been published. Mathys et al.^30^, used snRNA-seq to examine prefrontal cortex cell types including astrocytes. Although changes were observed, no explicit connection was made to A1 activation. We employed the ‘Gene Set Score’ method of Srinivasan et al.^31^ to compare astrocyte changes in one published AD data set, an ALS SOD1 mouse model^32^ and a human Parkinson’s disease (PD) data set^33^ with those changes observed in A-T. These results (Fig.4E) show a consistent directional correlation of astrocyte genes seen in A-T with the astrocyte gene sets derived from the Mathys AD studies. Additionally, the PD study showed a correlation between its upregulated genes and the changes seen in A-T.

Our NETSseq data revealed a striking down-regulation of synaptic transmission genes in A-T associated astrocytes. GSEA analysis showed that this downregulation is also observed for key genes in the Mathys et al.^30^ AD data set (Fig.S5B). This suggests that down-regulation of synaptic transmission genes is a common feature of activated astrocytes regardless of brain region or disease. Ongoing astrocytic dysfunction in regulating synaptic function may therefore be a phenomenon found across neurodegenerative diseases rather than being specific to A-T.

### Enrichment Analysis Reveals Accelerated Aging-Related Astrocytic Changes in A-T

Given the size and spread of age across the current cohort of control donors is relatively large, we were able to investigate the impact of normal aging on cerebellar gene changes and compare these findings with those from the A-T donors. PCA analysis of the most differentially expressed genes revealed that PC1 is associated with age in control donors. Interestingly, the subjects with A-T align with much older control donors, suggesting an accelerated aging phenomenon (Fig.4F). To better understand how A-T donors compare with age-matched controls, we analyzed the expression of A1-associated genes in younger and older control groups and A-T donors (Fig.4G). Interestingly, some markers showed similar expression levels between the two control groups but were upregulated in A-T donors, suggesting that neurotoxicity is more pronounced in the disease than in normal aging. In contrast, PAN markers such as GFAP, CD44, and CP were highly expressed in older controls compared to younger donors but showed an even greater elevation in expression in A-T donors (Fig. 4H), indicating an enhanced or accelerated activation signature of PAN-associated markers in A-T.

To further explore the changes occurring in donors with A-T and with aging, we performed enrichment analysis on the upregulated differentially expressed genes in both conditions (Table S4, Table S6). A consistent and significant enrichment of gene sets associated with changes in AD or associated with cognitive decline was observed (e.g. Mathys et al.,2019 - Astrocyte AD vs non-AD (up); Mostafavi et al., 2018 - ROSMAP-Clin AD enriched; Mostafavi et al., 2018- ROSMAP-Cognitive decline (UP)) in both aging and A-T. However, specific enrichment for gene sets associated with inflammation and astrocytic activation was seen in A-T only (Fig.4I, Table S7), further suggesting that the conversion to a more toxic astrocytic phenotype may be part of the normal aging process, but which is significantly accelerated as a consequence of the ATM mutated protein in A-T.

### Microglial A-T Changes Suggest an Accelerated Aging Phenotype

Lai et al.^13^ showed that activated microglia in the A-T cerebellum exhibit transcriptomic signatures resembling those seen in aging and neurodegeneration-associated microglia. However, they performed their analysis comparatively. Our microglia cohort consists of control donors with age ranges between 8 and 93 years old. Therefore, an assessment of the gene expression patterns associated with age were compared within the A-T dataset. PCA, based on the most variable genes identified through age regression analysis, demonstrated a significant correlation with age progression. Interestingly, donors with A-T are positioned at a similar level to older control donors (Fig.5A) suggesting that the microglial gene expression profiles seen in the A-T cerebellum mirrors the aging process but in much younger donors with A-T. This may suggest that the ATM mutation is accelerating the aging phenotype of microglia in A-T, contributing to an accelerated neurodegenerative disease process.

**Fig. 5.**
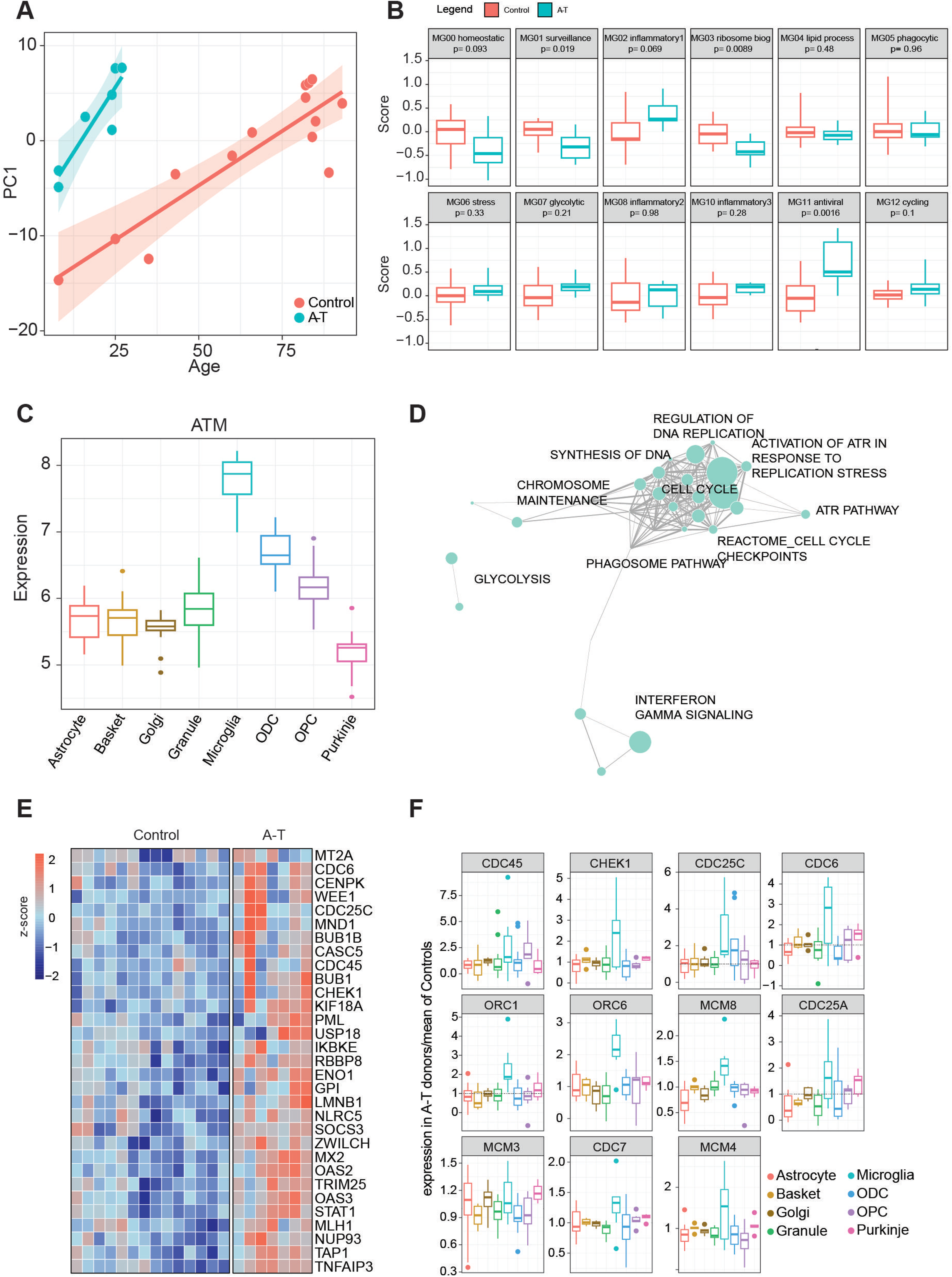
(A) Scatter plot between PC1 dimension and age, computed with the significant age-related differentially expressed genes in microglia. Linear regression analysis was split by disease condition. (B) Distribution of gene set scores for signatures identified in Sun et al. (C) Expression distribution of ATM gene in control donors across examined cell types. (D) KEGG pathway network enriched with the genes upregulated in Microglia and (E) heatmap of the significant differentially expressed genes (from D). (F) Normalised expression of A-T donors (over means of controls) of the GSEA leading-edge genes from “Activation of ATR in response to replication stress” indicates an A-T specific expression in microglia.

### Inflammatory Signatures in A-T: Implications for Neurodegenerative Pathways

Sun and colleagues^34^ identified 12 microglia clusters associated with distinct biological functions. Using gene set scoring to analyze our A-T microglia samples, we see that the homeostatic and surveillance functions, MG00 and MG01, are impaired in A-T while the inflammation pathways MG03, MG08 and MG10 are all elevated (Fig.5B). The homeostatic microglial cluster is crucial for maintaining normal brain function, while the surveillance cluster monitors and responds to the brain microenvironment for signs of cellular damage or infection. The downregulation of these clusters in A-T suggests a compromised ability of microglia to perform their normal essential homeostatic roles, potentially as a consequence of chronic inflammation and on going cellular stress which are hallmark features of A-T and other neurodegenerative conditions, such as AD. Interestingly the MG10 inflammation cluster was previously associated with changes in early AD, while the MG02 inflammation cluster was linked to late AD changes^34^. Overall, our A-T microglia signature reflects changes seen in late AD indicating a more advanced inflammatory state in A-T brains. Taking these microglial observations, together with the astrocyte inflammatory signaling changes, these findings reflect the importance of glial cell dysfunction in the pathophysiology of A-T, pointing towards disrupted immune surveillance and altered inflammatory responses across cell types as contributing factors to A-T disease progression.

### DNA Damage Response in A-T Microglia: Enrichment of ATR Regulation Highlights ATM Pathological Significance

A-T is characterized by defects in the ATM protein kinase^13^, which plays a central role in DNA damage response pathways. ATM is expressed at significantly higher levels in control donor microglia compared to the other cerebellar cell types examined here suggesting that microglia may be more affected by deficient ATM than other cell types (Fig.5C). This is likely attributable, at least in part, to the ability of adult microglia to proliferate as part of their normal homeostatic physiological condition and exacerbated during pathology-induced stimulation^35^.

Consistent with this hypothesis, the most upregulated pathway, as determined by the most overexpressed genes in A-T, is the Reactome gene set "Activation of ATR in response to replication stress" (Fig.5D). Within this group, genes involved in cell cycle checkpoint and DNA replication are consistently upregulated, including CHEK1 (Fig.5E), a positive regulator of cell cycle arrest. This potential adaptive response aligns with the importance of the ATR signaling in preserving genomic integrity under conditions of increased replication stress. This process is likely to be specific to glial cell types and particularly microglia during periods of elevated inflammation and cellular damage. Expression analysis of the leading-edge genes identified in the GSEA results associated with ATR response show a microglia-specific upregulation of these genes relative to other cell types (Fig.5F).

Among the 12 microglial clusters, MG11 was the most significant and differentially regulated between donors with A-T and control donors (Fig.5B). This gene set was associated with antiviral response, thus suggesting similarities in activation mechanisms needed for viral-induced responses and A-T microglia activation. It is known that cytosolic viral DNA induces type 1 interferon as a defence mechanism^36^. It has been argued previously that DNA damage or inhibited DNA repair can lead to nuclear DNA being transported into the cytoplasm leading to a ‘sterile inflammation response’ and the induction of interferons^36^. In support of this, genes in the KEGG pathway "Cytosolic DNA Sensing," which involves the detection of cytosolic DNA by immune receptors and triggers an antiviral response through the production of type I interferons and cytokines, are specifically expressed in microglia compared to other cerebellar cell types (Fig. S5C).

Collectively, these findings indicate that microglial pathways are significantly changed in A-T, and these disruptions may play a crucial role in driving aberrant neuroinflammation and progressive neurodegeneration^13,37^.

## Discussion

The comprehensive analysis of cerebellar post-mortem tissues from donors with A-T and control donors presented here has provided significant insights into the cell type specific changes occurring in A-T. Furthermore, the NETSseq platform achieved high-resolution nuclear RNA sequencing that outperformed single-nuclei RNA sequencing in detecting low-expressed and cell type specific genes and minimized cross-contamination among cell types. This enabled the identification of distinct gene expression patterns and changes in A-T, revealing significant disruptions to neuronal and glial cell function. Using deconvolution approaches we were able to accurately estimate the changes in cell type proportions from unsorted samples, with the same or higher precision than snRNA-seq studies.

We identified a significant loss of cerebellar granule and Purkinje neurons; the neuronal loss, coupled with changes in neurotransmitter signalling and synaptic function, highlights the disrupted motor coordination in A-T. Furthermore, our findings suggest a compensatory mechanism that increases ODC and OPC populations, possibly in response to ongoing myelin damage. However, it is clear from these data that glial subtypes likely have a significant role in this impaired neurotransmission and ongoing neurodegeneration and may even be a key trigger point. In addition to playing a significant role in shaping synaptic neurotransmission, cerebellar function and likely dysfunction, astrocytes together with microglia are also modulating ongoing inflammatory processes across multiple cell types, which may be driving or at least contributing to the progression of the disease process.

Astrocytes are widely recognized for their role in supporting neurons; they maintain homeostasis by regulating extracellular potassium concentrations and by removing neurotransmitters released at synapses, as well as controlling energy supply by regulating blood flow and providing lactate to neurons^38^. Moreover, neuron-astrocyte interactions and particularly the intimate interactions between astrocytic projections and synapses, in which astrocytes enwrap the pre- and post-synaptic neuronal elements, contribute to and regulate synaptic transmission^38^. This has been reported to be of particular importance for the BG-Purkinje neuron interactions within the cerebellum^29^ suggesting that the dysfunctions observed here may be more impactful in this particular brain structure, rendering the cerebellum more sensitive to disruption. In preclinical species these interactions have been shown to be involved in regulating behavioural function^39^ and complex information processing such as in cognition and memory processing^40,41^. Thus, many of the changes found in astrocytes (and neurons) may be contributing to a general impairment in synaptic function and support, leading to disrupted cerebellar signalling and neuronal communication. Our data further suggest that the neurotoxic and PAN signatures exhibit distinct expression patterns in astrocytes, with the neurotoxic signature more upregulated in A-T donors and changes in the PAN signature suggesting an accelerated aging phenomena in the cerebellum of A-T individuals.

Interestingly many of these glial functions also appear altered in other neurodegenerative disorders such as AD as well as normal aging; thus, the ATM mutation may be accelerating processes such as neuroinflammation in A-T. The microglia specific expression of the ATM gene and clear disruptions in the ATR pathway suggest that this cell type may be a central mediator of ongoing neuroinflammation. However, parallel changes in inflammatory markers in astrocytes and neurons indicate a complex interplay between cell types leading to aberrant inflammatory processes and the progressive loss of key neuronal networks. Further research is needed to fully understand these pathways, but these observations provide important insights into the pathophysiology of A-T.

In summary, this study provides a comprehensive cell type specific overview of gene expression and cellular composition in A-T cerebellum. NETSseq and these datasets provide a comprehensive high-resolution, cell type specific transcriptomic reference that can advance targeted investigation of disease mechanisms and therapeutic discovery in A-T.

## STAR Methods

### Human tissue

All human tissue donations used in this study were fully consented for research. Tissue was stored and data managed in compliance with the UK Human Tissue Act, with local Research Ethics Committee approval obtained from the Health Research Authority and in accordance with the World Medical Associations Declaration of Helsinki for medical research. Tissue was obtained from the NIH Neurobiobank at the University of Maryland, Baltimore, MD, Queen Square Brain Bank (The Queen Square Brain Bank is supported by the Reta Lila Weston Institute of Neurological Studies, UCL Queen Square Institute of Neurology), The Netherlands Brain Bank (Netherlands Institute for Neuroscience, Amsterdam); all material was collected from donors for or from whom a written informed consent for a brain autopsy and the use of the material and clinical information for research purposes had been obtained.

### Tissue preparation

Snap frozen cerebellar tissue samples were dissected on a brass plate, over dry ice. Tissue blocks for FFPE processing were first fixed in formalin at room temperature, the excess formalin drained and washed in water. Using an automated tissue processor (TP1020, Leica) the tissue was passed through a rising concentration of ethanol (70%, 80%, 95% and 100% twice for 1 h), then HistoChoice (twice for 1 h), before being immersed in Paraplast Plus (twice for 1 h). Tissues were placed in moulds, embedded in paraffin using a Paraffin Embedding Station (EG1150H, Leica), left to set on a cool plate (EG1150C, Leica), and were removed from the mould and stored at room temperature until required for sectioning. FFPE sections were cut from blocks at a thickness of 4 µm on a rotary microtome (Leica RM2255; 535 blades). Sections were floated in a water bath to flatten, before mounting on microscope slides. Slides were dried on a hotplate at 40°C for 4 h.

Frozen blocks were sectioned at a thickness of 10 µm in a freezing cryostat (Leica CM1850; MX35 blades), before mounting on microscope slides and stored at -80°C.

### Immunohistochemistry

Sections were fixed in 4% paraformaldehyde/phosphate buffered saline (PBS) for 10 min, washed, blocked for non-specific protein binding and endogenous peroxidase activity and then incubated with antibodies for 20 min at room temperature. Following primary incubation, sections were washed and incubated with horseradish peroxidase (HRP)-conjugated secondary antibodies (20 min, room temperature), washed again and then incubated with chromogen (10 min, room temperature). Sections were counterstained using hematoxylin, dehydrated through an ascending ethanol series, cleared and cover-slipped under Xylene.

Sections were evaluated using brightfield microscopy (Zeiss Axiophot) and IHC staining patterns were compared with negative control incubations (non-immune rabbit immunoglobulin G) performed under identical conditions on adjacent sections.

### *In situ* hybridization

RNAscope® 2.5 HD Assay (ACDBio) was carried out according to the manufacturer’s instructions, against the probes listed in Table S8, on cerebellum tissue from healthy donors. A GFAP probe was included as a positive assay control. Frozen sections were fixed with pre-chilled 4% paraformaldehyde for 15 min at room temperature and passed through a series of rising ethanol gradients (50%, 70% and 100% twice for 5 min each), then left at room temperature for 5 min to fully dehydrate. Sections were treated with hydrogen peroxide for 10 min, before washing in diethylpyrocarbonate-treated water for 5 min. Pre-warmed probes were applied for 2 h at 40°C, within a humidity-controlled chamber (HybEZ oven, ACD Bio). Following the incubation with probes, slides were washed twice in wash buffer (WB, ACDBio) for 2 min, at room temperature.

Signal amplification and detection reagents (RNAscope® 2.5 HD Detection Reagents-RED, ACDBio) were applied sequentially. Sections were incubated in AMP 1, AMP 2, AMP 3, and AMP 4 for 30, 15, 30 and 15 min, respectively, at 40°C, with 2 × 2 min WB washes in between. AMP 5, and AMP 6 reagents were applied for 30 and 15 min, respectively, at room temperature, with 2 × 2 min WB washes in between. Chromagen FAST RED was then applied for 10 min, at room temperature, before washing in WB twice for 2 min. The sections were counterstained with 50% Gill’s hematoxylin I (Pioneer Research Chemicals) for 30 s at room temperature, rinsed, and dried for 15 min at 60°C, before applying a coverslip with VectaMount (Vector Laboratories).

### Quantitative histopathology

Sections immunohistochemically stained using a polyclonal antibody to RBFOX3 were digitally scanned (Hamamatsu Nanozoomer). Three neuroanatomically-matched fields of view were selected from three normal donors and four donors with A-T. Image processing and analysis was carried out using Image J (Ver.1.51 National Institutes of Health). Positively stained nuclei were accurately segmented according to both colour and staining intensity and automatically counted. Adjustments based on the average size of positive nuclei were made for aggregates.

### Nuclei preparation

Nuclei isolation protocols described fully in previous publications^11^, were miniaturized to enable a more efficient and higher throughput method to process larger donor numbers.

To isolate nuclei, cerebellar frozen tissues were thawed on ice for a minimum of 30 min and transferred to 1 mL of homogenization medium (0.25 M sucrose, 150 mM KCl, 5 mM MgCl_2_, 20 mM Tricine pH 7.8, 0.15 mM spermine, 0.5 mM spermidine, EDTA-free protease inhibitor cocktail, 1 mM DTT, 20 U/mL Superase-In RNase inhibitor, 40 U/mL RNasin ribonuclease inhibitor). Using a 2 mL glass dounce tissue grinder set, samples were homogenized by 30 strokes of a large clearance pestle A (0.003-0.005 in.) followed by 30 strokes of small clearance pestle B (0.005 – 0.0025 in.). Homogenate was adjusted to 1.92 mL of homogenization medium and supplemented with 1.78 mL of a 50% iodixanol solution (50% Iodixanol/Optiprep, 150 mM KCl, 5 mM MgCl_2_, 20 mM Tricine pH 7.8, 0.15 mM spermine, 0.5 mM spermidine, EDTA-free protease inhibitor cocktail, 1 mM DTT, 20 U/mL Superase-In RNase inhibitor, 40 U/mL RNasin ribonuclease inhibitor), and laid on a 27% iodixanol cushion. Nuclei were pelleted by centrifugation for 25 min, 10,000 rcf, 4°C in an Eppendorf 5427 R centrifuge, FA-45-12-17 rotor. The nuclear pellet was resuspended in homogenization buffer.

### Nuclei labelling and sorting

After isolation, resuspended nuclei were fixed with 1% formaldehyde for 8 min at room temperature, and then quenched with 0.125 M glycine for 5 min. Nuclei were pelleted at 1000 rcf, 4 min, 4°C in a FA-45-30-11 rotor, and then washed once with homogenization buffer and once with Wash Buffer (PBS, 0.05% TritonX-100, 50 ng/mL BSA, 1 mM DTT, 10 U/µL Superase-In RNase Inhibitor). Nuclei were blocked with Block Buffer (Wash buffer with an additional 50 ng/mL BSA) for 30 min, incubated with primary/fluorescently conjugated antibody for 1 h, and then washed three times with Wash Buffer with spins in between washes as described above. Nuclei were then incubated in secondary antibody for 30 min and washed three times with Wash Buffer. All incubation steps were performed at room temperature. Primary, secondary and conjugated antibodies were diluted in Block Buffer.

### Primary/conjugated antibodies

Antibody concentrations were either validated previously^11^ or determined by evaluation of antibodies in validation tissue by flow cytometry (Table S8).

### Secondary antibodies

Secondary antibodies were prepared according to manufacturer’s instructions (Jackson ImmunoResearch) for extended storage and rehydration by adding an equal volume of glycerol (final concentration of 50%) and used at a 1:250 dilution (Table S8).

### Flow cytometry

Prior to flow cytometry, nuclei were co-stained with DAPI to 0.01 mg/mL final concentration. Nuclei were analyzed and sorted using a BD FACSAria Fusion (BD Biosciences, San Jose, CA, USA) flow cytometer using the 355 nm, 488 nm, 561 nm, and 640 nm lasers. All samples were first gated using an FSC/SSC gate to exclude debris, followed by a single nuclei DAPI gate to exclude aggregated nuclei. Analysis was performed using FACSDiva (BD) or FlowJo software. To reduce RNA degradation all nuclei samples were sorted at 4°C and stored at -80°C, before being processed for RNA extraction.

Two different antibody panels were used to isolate cell types. RBFOX3, FOXK2, ITPR1, OLIG2 and IRF8 were used to sort Purkinje cells, basket cells, granule cells, Golgi cells, oligodendrocytes, oligodendrocyte precursor cells and microglia. RBFOX3, SLC1A3, OLIG2 and IRF8 were used to sort oligodendrocytes, oligodendrocyte precursor cells, microglia and astrocytes. For an exhaustive list of antibodies used refer to Table S8. The Mann–Whitney U test was used to check for significant differences in percentage cell-type composition between control and AT samples for astrocytes, oligodendrocytes and oligodendrocyte precursors. Linear regression was used to rule out age as an explanatory variable.

### RNA extraction and library builds

RNA extractions from sorted nuclei were carried out with FFPE RNA Purification kits (Norgen: Product # 25300) using the manufacturer’s protocol from Step 2 (Lysate Preparation) with the following modifications: The On-column DNA removal Protocol was omitted. RNA binding and Column Wash centrifuge steps were carried out at 1,789×g for 2 min, followed by 1,789×g for 5 min to dry the resin. Elution was carried out in two steps, with 25 µL Elution Solution A added to each column followed by a 1 min room temperature incubation and centrifugation at 200×g for 2 min, then a further 15 µL Elution Solution A was added to each column with centrifugation at 1,789×g for 5 min. For samples containing >2,000 nuclei, the final 30 µL elution was split into two 15 µL aliquots. Purified RNA was vacuum dried in a CentriVap Benchtop Vacuum Concentrator (Labconco) for 40 min (or until no liquid was visible) at 4°C and either used directly for library preparation or stored at -20°C.

Sequencing library generation was carried out using the Trio RNA-Seq, Human rRNA Library Preparation Kit (Tecan: Product # 0506-A01) using the manufacturer’s protocol with the following modifications: Lyophilized RNA was resuspended in 12.5 µL DNAse mix and incubated at 37°C for 30 min followed by 60°C for 5 min and 4°C hold. Library amplification I was carried out as 72°C 2 min/95°C 2 min/2×(95°C 30 s/ 60°C 60 s)/6×(95°C 30 s/ 65°C 60 s)/65°C 5 min/4°C hold. Library Amplification II was carried out as 95°C 2 min/2×(95°C 30 s/ 60°C 60 s)/6×(95°C 30 s/ 65°C 60 s)/65°C 5 min/4°C hold. Final libraries eluted in 30 µL DNA Resuspension Buffer (DR1) were checked on Agilent Bioanalyzer DNA 1000 chips, then measured on a Nanodrop 8000. 32 sample pools (normalized to 500 ng/sample then diluted to 5 ng/µL) were sent for sequencing on an Illumina NextSeq High Output platform.

### Bioinformatics analysis

FastQC (http://www.bioinformatics.babraham.ac.uk/projects/fastqc/) was used for initial quality control. Adapter trimming was performed with Cutadapt v1.12^42^. Reads were aligned to the hg38 human genome build (NCBI primary analysis set) using STAR aligner^43^ v2.5.2b with splice junction files generated from Gencode release 27 and RefSeq GRCh38.p10. Post-alignment quality control was performed by examination of the STAR alignment metrics, Picard tools v2.8.2 (CollectRNASeqMetrics) (https://broadinstitute.github.io/picard/) and RSeQC v2.6.4 (tin.py)^44^. Expression quantitation was performed with htseq-count v0.6.1p1^45^ using a custom annotation file based on RefSeq where both exonic and intronic reads are counted for each gene.

The read counts table from each sample was imported into RStudio (R version 4.0.5) running under Red Hat Enterprise Linux 8. Using edgeR^46^, the read counts were normalized into Counts Per Million (TPM), log transformed, and batch corrected. For each individual cell type, we removed technical variance associated with batch effect using the sva R package^47^. Briefly, the sva R package provides detection of unwanted variation in high-dimensional data. This unwanted variation, often caused by batch effects, technical artifacts, or unknown confounding factors, can obscure the true biological signals of interest. We incorporated relevant surrogate variables in the differential analysis models; the models aim to assess differences due to the disease condition (or any variable of interest such as age), while accounting for covariates or potential confounders. Differentially expressed genes were identified using DESeq2 R package^48^.

Hierarchal clustering analysis was performed using the pheatmap R package^49^. Principal component analysis was performed using the pcaMethods R package^50^. Metabolic flux analysis was computed using the scFEA R package^51^. Briefly, scFEA is a tool designed to infer metabolic activity based on gene expression data. It works by modelling the flux of metabolites through predefined metabolic modules, which represent groups of reactions within metabolic pathways. Using transcriptomic data as input, scFEA predicts the activity of these modules by integrating gene expression levels with a flux balance analysis framework. This allows for the estimation of pathway activity across the samples, providing insights into cellular metabolism and its variation across different conditions. Cell type specific genes were identified using the specificity index algorithm previously described in^11^ and^52^. log_2_(TPM) values were used as gene expression input and ranks were averaged across 1,000 iterations.

### Quantification of cell type proportions

To quantify cell type proportions in the unsorted populations we performed a deconvolution analysis using the deconRNAseq R package^53^. deconRNAseq works by leveraging reference expression profiles, which are known RNA expression levels of different cell types, to analyze the mixed tissue sample’s RNA sequencing data. It uses these profiles to construct a mathematical model that deconvolutes the mixture, identifying the contribution of each cell type. The algorithm optimizes the fit between the observed mixed sample data and a linear combination of the reference profiles, thereby estimating the proportion of each cell type present in the sample. We assembled the reference dataset using our sorted, cell type specific, RNA-seq expression profiles. This method enabled us to estimate the relative abundances of each cell type in the unsorted samples. For the comparative deconvolution analysis, we first created a reference panel for each cell type. Each reference panel consisted of two expression signatures: one representing the target cell population and the second representing the average gene expression of all other cell types. This resulted in nine distinct signature datasets, each corresponding to a different cell type in comparison to the rest. Next, we compared the gene expression profiles of the unsorted samples against each of these reference signatures. For each comparison, we quantified the fraction of similarity between the unsorted sample and a given target cell type, relative to the expression of all other cell types. This approach allows us to assess the proportion of similarity between the unsorted samples and individual cell types, providing a more accurate means of identifying changes in the proportion of specific cell types between experimental conditions.

### Gene Set Enrichment Analysis (GSEA) and gene set scores

Gene Set Enrichment Analysis (GSEA) was performed using the method described by Subramanian et. al.^54^ on gene sets generated from the Molecular Signatures Database (MSigDB) v6.2 against a ranked list of genes derived from the DESeq2 analysis results. Pre-ranked gene lists (.rnk files) were created from the DESeq2 Wald statistic, low expressed genes with baseMean less than 10 were filtered out to reduce noise.

Microglial state marker genes were harvested from^34^ as is. For each state, labelled MG0∼MG12, genes significantly upregulated (adjusted P-value<0.05) and with a minimum fraction of cells expressing the gene of 0.25 were used to build the gene sets for GSEA.

Similarly, we harvested disease-related and cell type specific gene sets from several relevant publications. These gene sets represent up- or down-regulation with respect to key Alzheimer’s disease traits. We extracted significantly differentially expressed genes (adjusted P-value<0.05) and subdivided them based on the direction of their fold changes.

To compare changes between gene expression profiles we generated with other changes seen in other neurodegenerative diseases or models we used the gene set score method described in^31^.

## Acknowledgments

We would like to thank the donors and their families for the generous contribution of the tissues used in the present study. We would like to thank the A-T Children’s Project for their help in sourcing the A-T tissue and guidance during these studies. We would like to thank Professor Nathaniel Heintz for his invaluable input to the research that went into these studies and the current manuscript.

## Supplementary Figures

**Fig. S1.**
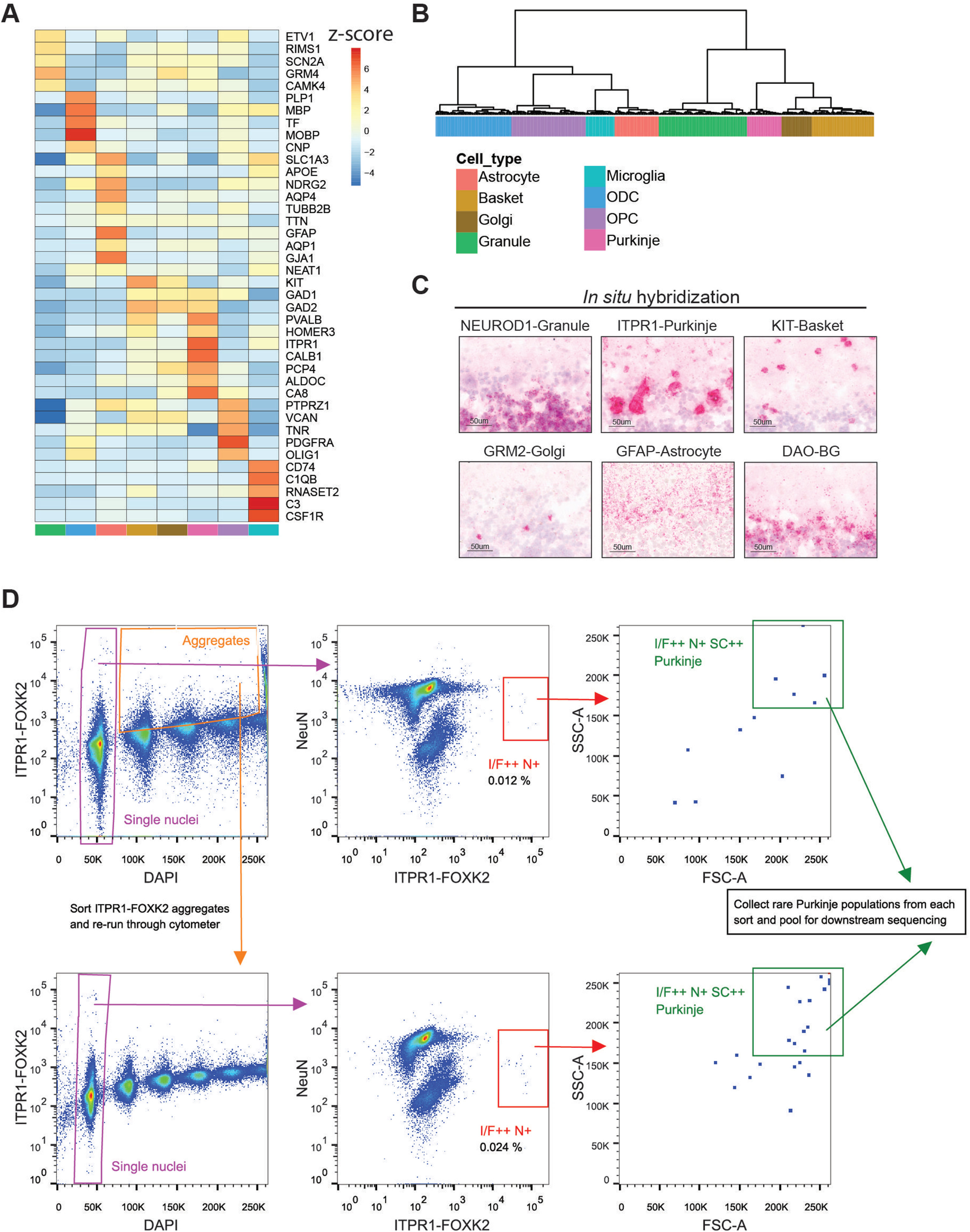
(A) Heatmap of cell type specific genes in each cerebellum cell type (markers from^13^). (B) Dendrogram of cerebellar cell types in our study computed with all the expressed genes (scaled expression of all genes and Ward2 clustering were used). (C) In situ hybridization (ISH) showing characteristic cell type specific identity marker genes chosen from NETSseq data in different cell types of the cerebellum. Labelled cells appear distinct in their expected locations. Sections show the molecular layer at the top, granule cell layer beneath and the Purkinje layer at the transition. (D) Representative example of the gating strategy for Purkinje nuclei in human cerebellar tissue. First, single nuclei (purple gate) are identified by DAPI. Single nuclei are visualised by ITPR1-FOXK2 vs NeuN, where the I/F++N+ (ITPR1/FOXK2++, NeuN+) population is identified (red gate). To further exclude contamination from smaller and less complex cell types, the I/F++N+ population is refined using FSC-A and SSC-A to give the I/F++N+SC++ population (green gate) which is the targeted Purkinje nuclei. Due to the very low abundance of Purkinje nuclei, we also re-sort ITPR1-FOXK2 positive aggregates (orange gate) and follow the same gating strategy as just described. This process of re-sorting aggregates can be repeated multiple times and all I/F++N+SC++ populations pooled to increase total nuclei number for downstream sequencing.

**Fig. S2.**
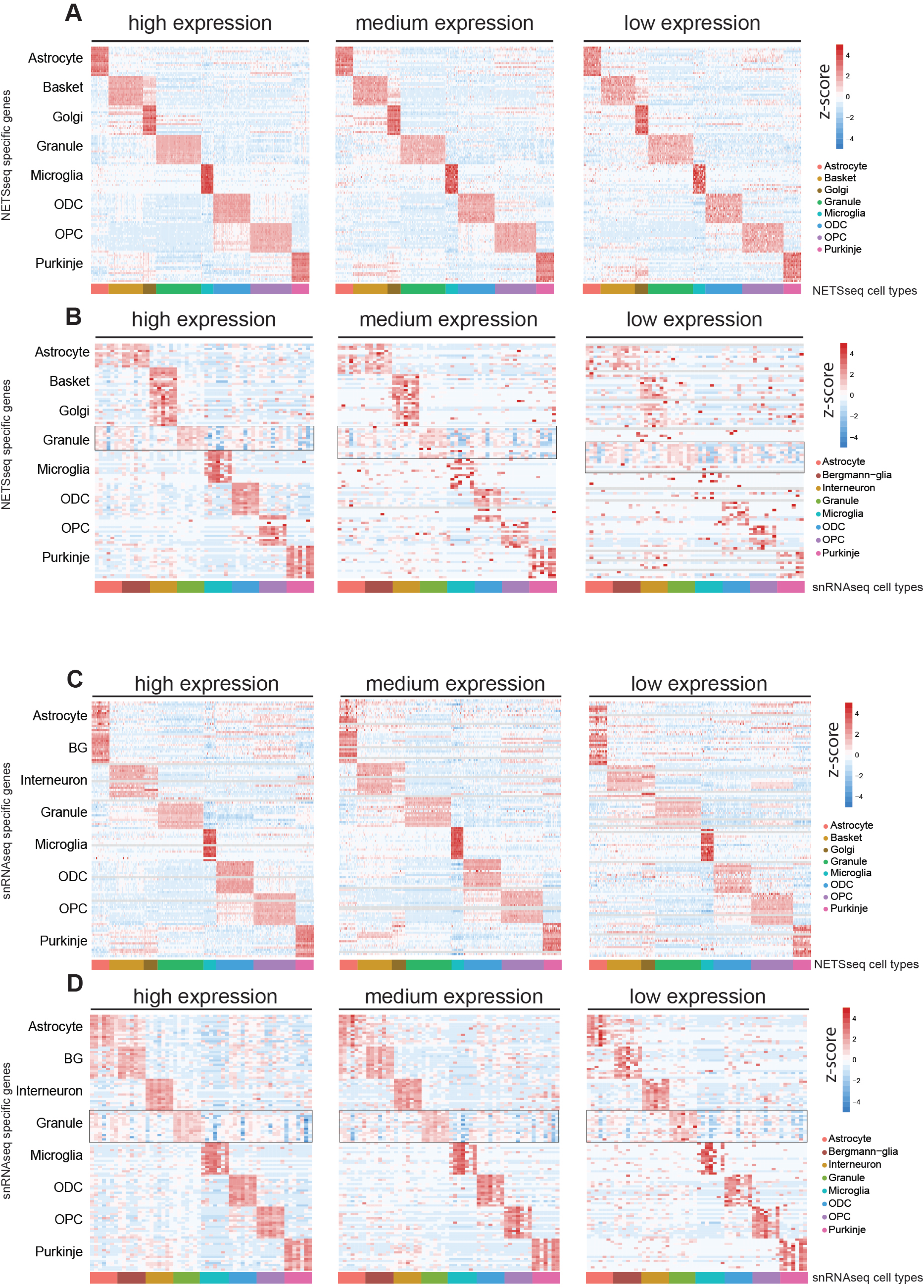
(A) Heatmap of NETSseq gene expression for cell type specific genes identified from the NETSseq cohort. Expression of the NETSseq cohort was split by quantile expression and clustered by cell type. The Y axis shows the cell type origin of the genes. The X axis shows the samples grouped by cell type. The specific signal was stable across the different expression thresholds. (B) Heatmap of snRNA-seq gene expression pseudo-bulked by donors for cell type specific genes identified from the NETSseq cohort, organised as above. The specificity signal was disrupted and lost for lower expressed genes. The black rectangle highlights granule contamination across different cell types, expression of granule specific genes was found across the dataset. (C) Heatmap of NETSseq gene expression for cell type specific genes identified from the snRNA-seq cohort, organised as above. The specificity signal was consistent across the different expression threshold, indicating that NETSseq platform doesn’t suffer from the same detection limits. (D) Heatmap of snRNA-seq gene expression pseudo-bulked by donors for cell type specific genes identified from the snRNA-seq cohort, organised as above. The square highlights granule contamination across different cell types even when the specific genes were identified using the snRNA-seq cohort.

**Fig. S3.**
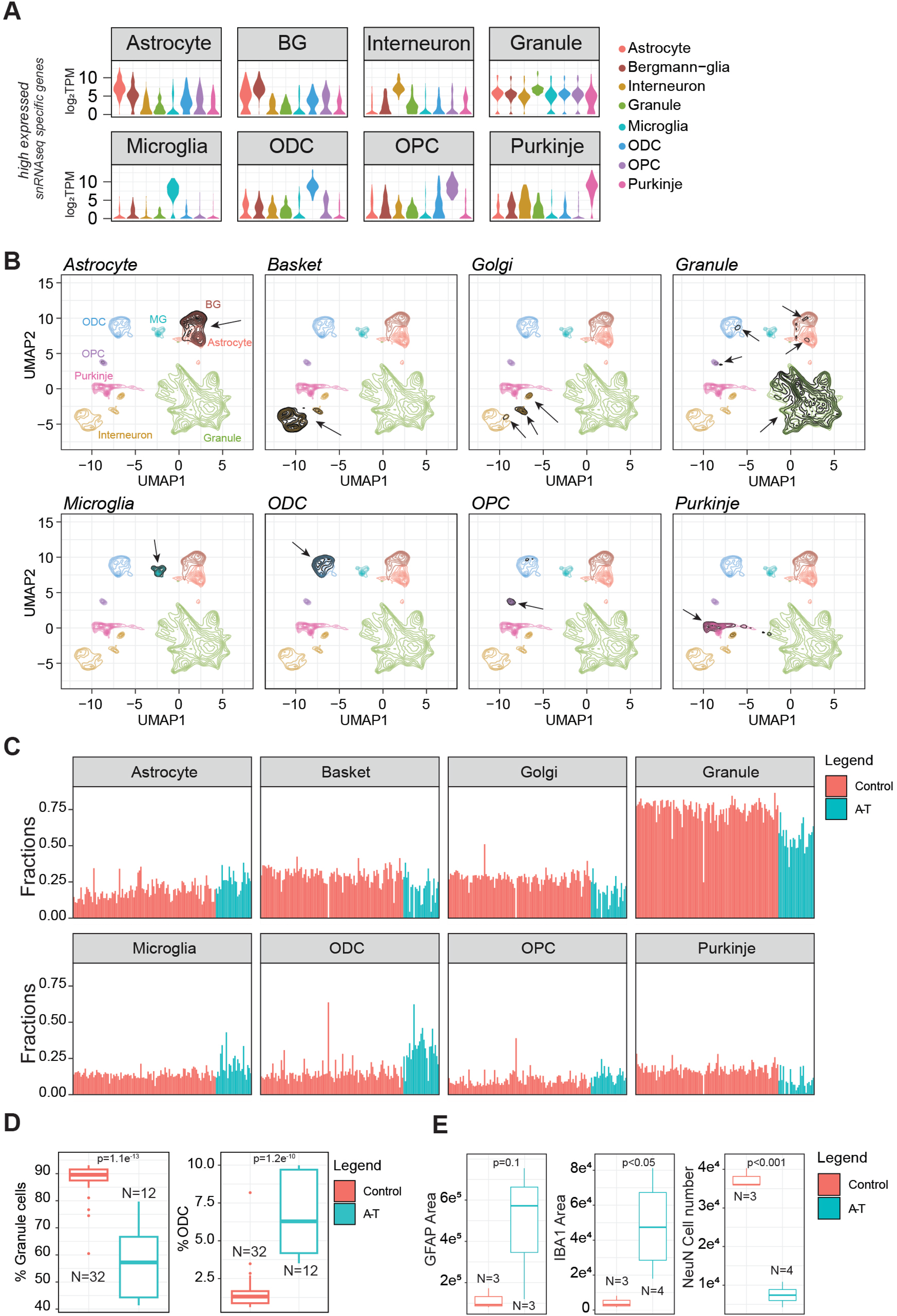
(A) Violin plot for the most expressed and cell type specific genes identified in the snRNA-seq cohort and their expression across the different cell types. Each panel shows one set of cell type specific genes (identified in the panel heading) expressed across all eight cell types. The granule panel highlights the level of granule contamination across the different cell types. (B) Density plot for the single nuclei dataset colored by the assigned cell types identified in the snRNA-seq study. The black density represents the expression density of the cell type specific marker identified in NETSseq cohort for the most expressed genes. The expression density overlaps entirely with the assigned cell types, indicating that the gating strategy used in our NETSseq platform isolated the correct population of cells and not subtypes. Black arrows indicate the expression density for the relevant cell type-specific genes. (C) Absolute fraction of similarity for each unsorted sample, coloured by disease condition. (D) Quantification of sorted nuclei from FACS for the specified cell types. (E) Quantitative image analysis of neurons and glia markers in cerebellum.

**Fig. S4.**
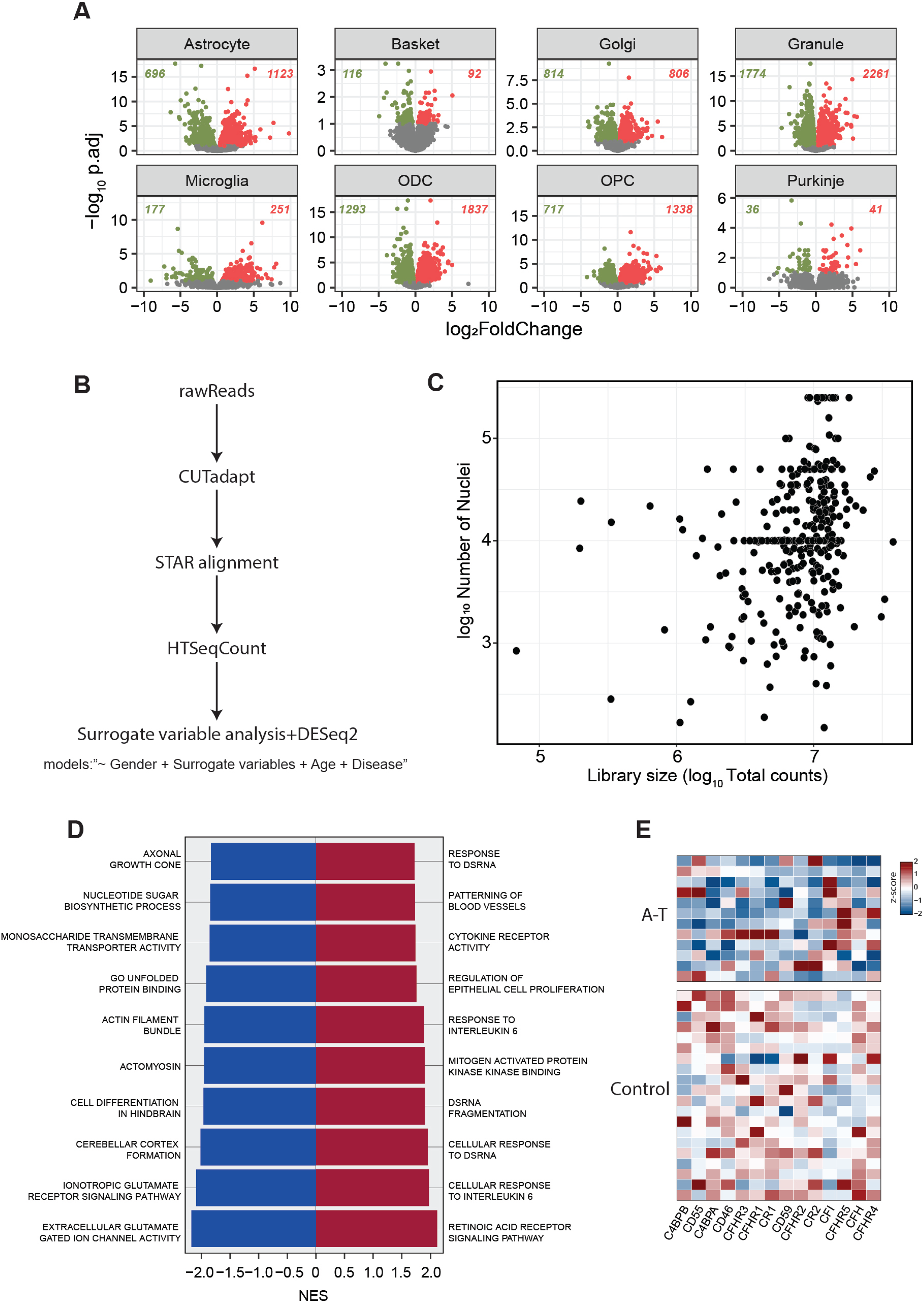
(A) Volcano plot depicting the differentially expressed genes between A-T and controls across cell types, coloured dots show the significant up or downregulated genes with the relative annotation (-log_10_p.adj>1 and baseMean>10). (B) Schematic representation of the pipeline used to compute differentially expressed genes. (C) Scatter plot showing the number of nuclei and the library size for each sequenced sample in our cohort. (D) Top 10 up and downregulated enriched gene ontology processes found in Purkinje neurons (pval < 0.01). (E) Inhibitors of complement components and their expression in A-T and control samples in astrocytes.

**Fig. S5.**
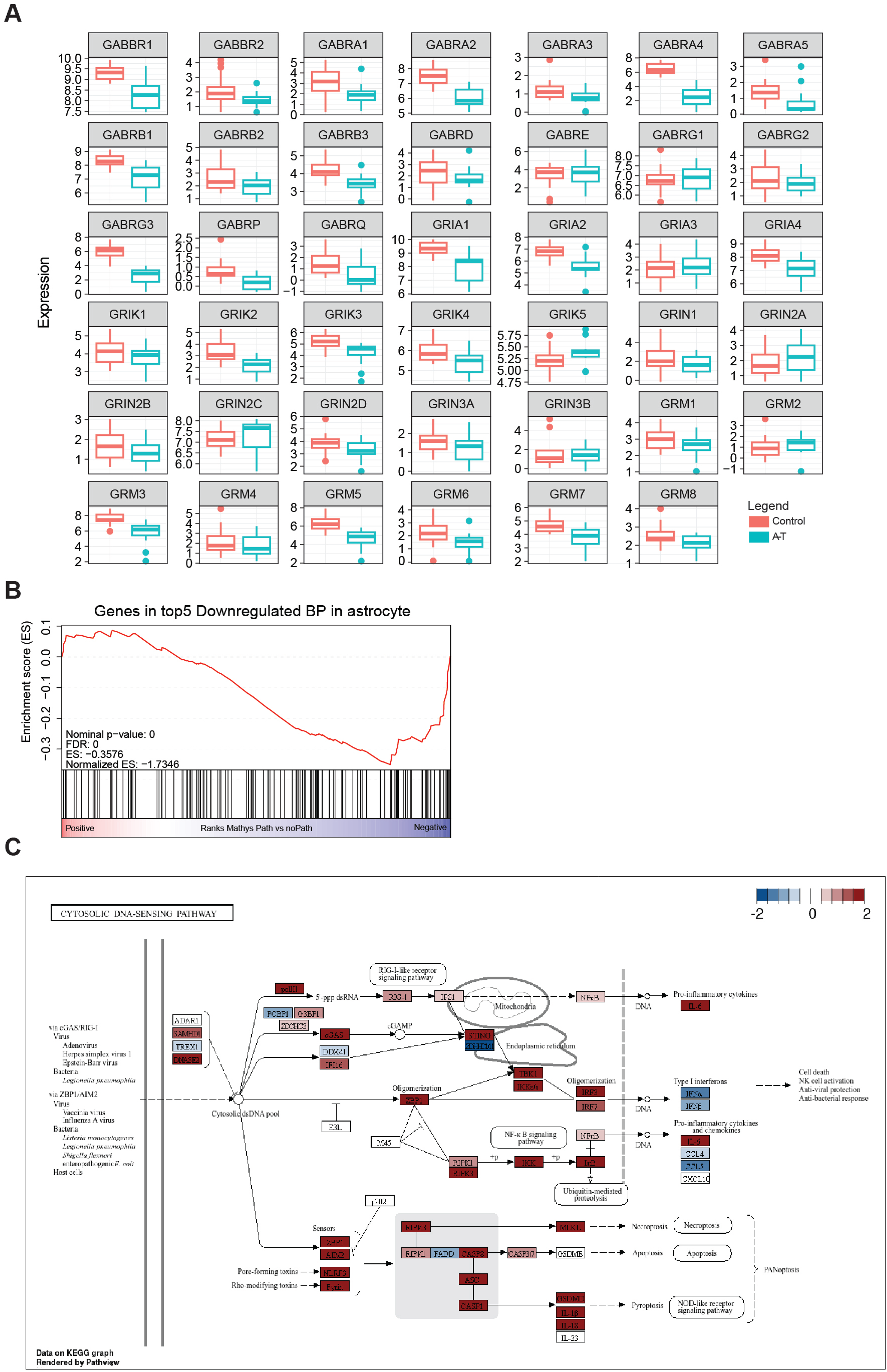
(A) Expression plot of neurotransmitters and receptors in A-T and control astrocytes. (B) Gene set score for the genes in top 5 BP from Fig.4E (regulation of synaptic transmission glutamatergic, neuron-neuron synaptic transmission, presynaptic process involved in synaptic transmission, regulation of postsynaptic membrane potential and gamma aminobutyric acid signaling pathway). (C) KEGG pathway “Cytosolic DNA sensing”. Genes are coloured by z-score value in microglia computed across all examined cell types (red indicates genes more expressed in microglia compared to other cell types).

